# Genome variation and meiotic recombination in *Plasmodium falciparum*: insights from deep sequencing of genetic crosses

**DOI:** 10.1101/024182

**Authors:** Alistair Miles, Zamin Iqbal, Paul Vauterin, Richard Pearson, Susana Campino, Michel Theron, Kelda Gould, Daniel Mead, Eleanor Drury, John O’Brien, Valentin Ruano Rubio, Bronwyn MacInnis, Jonathan Mwangi, Upeka Samarakoon, Lisa Ranford-Cartwright, Michael Ferdig, Karen Hayton, Xinzhuan Su, Thomas Wellems, Julian Rayner, Gil McVean, Dominic Kwiatkowski

**Affiliations:** MRC Centre for Genomics and Global Health, University of Oxford, Oxford, UK; Malaria Programme, Wellcome Trust Sanger Institute, Hinxton, UK; Wellcome Trust Centre for Human Genetics, University of Oxford, Oxford, UK; Bowdoin College, Brunswick, Maine, USA; Broad Institute of Harvard and MIT, Cambridge, Massachusetts, USA; Department of Biochemistry, Medical School, Mount Kenya University, General Kago Road, Thika, Kenya; Institute of Infection, Immunity and Inflammation, College of Medical, Veterinary and Life Sciences, University of Glasgow, Glasgow, UK; Eck Institute for Global Health, Department of Biological Sciences, University of Notre Dame, Notre Dame, Indiana, USA; Laboratory of Malaria and Vector Research, National Institute of Allergy and Infectious Disease, National Institutes of Health, Bethesda, Maryland, USA; Department of Statistics, University of Oxford, Oxford, UK

**Keywords:** malaria, plasmodium falciparum, next-generation sequencing, genome variation, INDEL variation, complex variation, meiotic recombination, gene conversion, copy number variation, drug resistance

## Abstract

The malaria parasite *Plasmodium falciparum* has a great capacity for evolutionary adaptation to evade host immunity and develop drug resistance. Current understanding of parasite evolution is impeded by the fact that a large fraction of the genome is either highly repetitive or highly variable, and thus difficult to analyse using short read technologies. Here we describe a resource of deep sequencing data on parents and progeny from genetic crosses, which has enabled us to perform the first genome-wide, integrated analysis of SNP, INDEL and complex polymorphisms, using Mendelian error rates as an indicator of genotypic accuracy. These data reveal that INDELs are exceptionally abundant, being more common than SNPs and thus the dominant mode of polymorphism within the core genome. We use the high density of SNP and INDEL markers to analyse patterns of meiotic recombination, confirming a high rate of crossover events, and providing the first estimates for the rate of non-crossover events and the length of conversion tracts. We observe several instances of recombination that modify copy number variants associated with drug resistance, demonstrating a mechanism whereby fitness costs associated with resistance mutations could be compensated and greater phenotypic plasticity could be acquired. We describe a novel web application that allows these data to be explored in detail.

## Introduction

Genome variation in the eukaryotic pathogen *Plasmodium falciparum* underpins both fundamental biology, such as the ability of the parasite to evade the human immune response, and clinical outcomes, through the evolution of antimalarial drug resistance. This is of particular concern with the recent spread of resistance to front-line therapies in South-East Asia (Ashley et al. 2014).

High throughput sequencing is a proven technology for the study of genome variation in *P. falciparum* and has yielded insights into natural patterns of variation and population structure (Manske, Miotto et al. 2012; Miotto et al. 2013), the generation of antigenic diversity (Claessens et al. 2014) and the genetic basis for artemisinin resistance (Ariey et al. 2014). Despite these recent advances, our current understanding of *P. falciparum* genome variation remains incomplete due to multiple factors that are challenging both for the sequencing technology and for the analytical methods used for variant discovery and genotyping. The highly compact 23Mbp genome has an extremely biased nucleotide composition, with 80.6% (A+T) content overall and ~90% (A+T) in noncoding regions (Gardner et al. 2002). As a result, many regions of the parasite genome are highly repetitive, with short tandem repeats and other low complexity sequences unusually abundant in both coding and non-coding regions (Gardner et al. 2002; Zilversmit et al. 2010; Muralidharan and Goldberg 2013; DePristo et al. 2006). Another difficulty is that parasite genes encoding antigenic targets of the host immune system tend to exhibit very high levels of diversity, such that individual alleles are typically highly diverged from the reference sequence. An extreme example of this phenomenon is the multi-copy *var* gene family encoding erythrocyte surface antigens which can diversify within the course of a single infection by non-allelic recombination (Bopp et al. 2013; Freitas-Junior et al. 2000; Claessens et al. 2014).

These factors have limited progress and there are a number of current knowledge gaps. There has been no comprehensive survey of insertion/deletion (INDEL) variation in *P. falciparum,* although there is evidence that INDELs may be unusually abundant (Jeffares et al. 2007; Haerty and Golding 2011; Tan et al. 2010). Little is known about variation in non-coding regions, which could have a significant impact on phenotype by regulating gene expression (Mok et al. 2014; Gonzales et al. 2008). Knowledge of complex variation, where haplotypes are highly diverged from the reference genome, is constrained to a few well-studied genes such as *msp1* (Roy et al. 2008).

A critical step in overcoming these obstacles is to have a reliable, empirical indicator of genotyping error, which allows different genotyping methods to be calibrated and compared. There are many potential sources of error in the process of high throughput sequencing and variant calling (Robasky et al. 2014) and different analytical methods may have different strengths and weaknesses. A proven approach is to sequence multiple individuals belonging to a pedigree and to identify genotype calls that are in violation of Mendelian inheritance. A small number of Mendelian inconsistencies are expected due to *de novo* mutation, but the observation of many inconsistencies is a strong indicator of genotyping error. Mendelian errors can thus be used to calibrate methods and filter data (Saunders et al. 2007).

Here we report an analysis of *P. falciparum* genome variation in the parents and progeny of experimental genetic crosses. We have studied all three crosses that have been published to date, involving the parental clones 3D7, HB3, Dd2, 7G8 and GB4, representing a broad range of genetic and phenotypic diversity (Su et al. 2007; Ranford-Cartwright and Mwangi 2012). It is worth noting that these genetic crosses were complex and laborious to produce, involving passage through blood cultures, mosquitoes (where sexual reproduction takes place) and non-human primates (Walliker et al. 1987). They have led to key discoveries regarding the causes of drug resistance (Wellems et al. 1991) and host specificity (Hayton et al. 2008). Although only a limited number of crosses are currently available, typically more than 30 genetically distinct progeny clones can be obtained from a single cross (Ranford-Cartwright and Mwangi 2012). The large number of progeny provides a higher power to observe Mendelian errors than smaller pedigrees or trios, and thus to identify variants which are spurious or where genotyping is unreliable. These three crosses are therefore a precious resource for studying genome variation in *P. falciparum,* because they represent the only experimental system within which Mendelian errors can be observed and used to guide variant discovery.

We use a combination of methods for variant discovery, leveraging both alignment of sequence reads to the 3D7 reference genome (DePristo et al. 2011; Li and Durbin 2009; McKenna et al. 2010) and reference-free sequence assembly (Iqbal et al. 2012) to build a map of genome variation within each cross incorporating SNP, INDEL and complex polymorphisms, and spanning both coding and non-coding regions. All of the variants included in the final call sets are highly consistent with Mendelian inheritance and show almost perfect genotype concordance between biological replicates. These data reveal some unique features of genome variation in *P. falciparum,* including an exceptionally high abundance of INDELs relative to SNPs. Because these data are a rich resource and valuable for further research in the genome biology of eukaryotic pathogens, we describe a novel web application providing a means for exploring and interacting with the data in an intuitive way.

We also address open questions regarding meiotic recombination in *P. falciparum,* a key biological process that generates and maintains genetic diversity in natural parasite populations, and thus contributes to parasite evolution. Previous studies using lower-resolution genotyping methods have estimated crossover (CO) recombination rates (Su et al. 1999; Jiang et al. 2011; Walker-Jonah et al. 1992) and provided evidence for non-crossover (NCO) recombination (Su et al. 1999). One study used high-throughput sequencing to resolve recombination events in two progeny of the HB3xDd2 cross, finding evidence that rates of CO and NCO recombination may be similar (Samarakoon et al. 2011b). Previous studies have also demonstrated that at least some recombination events occur within coding regions (Kerr et al. 1994) and suggested that recombination events are not uniformly distributed over the genome (Jiang et al. 2011). Here we combine SNP and INDEL markers to obtain a resolution of ~300bp within each cross, which is sufficient to differentiate between crossovers and non-crossovers, to estimate rates for both types of recombination event, and to study conversion tract lengths. This resolution is also sufficient to resolve the location of most recombination events relative to gene and exon boundaries and study the rate of intragenic recombination.

Finally we investigate recombination in the context of two large regions of copy number amplification, both of which segregate in the crosses and are associated with drug resistance (Wellems et al. 1990; Samarakoon et al. 2011a). We find evidence for crossover recombination within these amplifications leading to regions of pseudo-heterozygosity within progeny clones. This has potential implications for the evolution of drug resistance, because it demonstrates a mechanism whereby parasites could acquire both wild-type and mutant alleles at a drug resistance locus, which in turn could compensate for fitness costs associated with drug resistance mutations and create flexibility to adapt to variable drug pressure.

## Results

### Whole genome sequencing and genome accessibility

Whole genomes of parent and progeny clones from the crosses 3D7xHB3 (Walliker et al. 1987), HB3xDd2 (Wellems et al. 1990, 1991) and 7G8xGB4 (Hayton et al. 2008) were sequenced using Illumina high throughput technology, with the majority of samples obtaining an average depth above 100X (Table 1; Table S1). All DNA libraries were derived from haploid parasite clones in culture, and sufficient DNA was available to use PCR-free library preparation throughout, which has been shown to reduce some of the biases associated with the AT-rich *P. falciparum* genome and hence improve the evenness of coverage across both coding and non-coding regions (Kozarewa et al. 2009). PCR can also induce false INDELs, providing an additional motivation for using PCR-free libraries (Fang et al. 2014). The clone HB3 is a parent in two crosses, however because DNA samples were obtained from different sources and had different culturing histories the two HB3 clones were sequenced and genotyped separately, and are here labelled HB3(1) and HB3(2)

**Table 1.** Summary of sequence and variation data generated in this study for the three crosses 3D7xHB3, HB3xDd2 and 7G8xGB4. ^1^Number of independent recombinant progeny that were sequenced as part of this study and yielded usable sequence data. ^2^Coverage for each sample was calculated as the mean depth of sequenced bases across the whole genome. Values shown are the median (minimum-maximum) values of sample coverage within a cross. ^3^Total number of segregating SNPs combined from both alignment and assembly calling methods that passed all quality filters. ^4^Total number of segregating INDELs combined from both alignment and assembly calling methods that passed all quality filters. ^5^Nucleotide diversity (number of segregating SNPs per kilo base pair) calculated over 10kb non-overlapping windows within the core genome. Values shown are the median (5-95th percentile). ^6^Indel diversity (number of segregating INDELs per kilo base pair) calculated over 10kb non-overlapping windows within the core genome. Values shown are the median (5-95th percentile). ^7^Average distance between combined SNP and INDEL markers. Values shown are median (5-95th percentile).

|  | 3D7xHB3 | HB3xDd2 | 7G8xGB4 |
| --- | --- | --- | --- |
| No. progeny<sup>1</sup> | 15 | 35 | 28 |
| Coverage<sup>2</sup> | 102X (41-173) | 110X (22-637) | 107X (55-250) |
| No. SNPs<sup>3</sup> | 15388 | 14885 | 14392 |
| No. INDELs<sup>4</sup> | 26699 | 21576 | 20079 |
| Nucleotide diversity (kbp<sup>-1</sup>)<sup>5</sup> | 0.5 (0.1-1.6) | 0.5 (0.1-1.4) | 0.5 (0.1-1.4) |
| INDEL diversity (kbp<sup>-1</sup>)<sup>6</sup> | 1.3 (0.5-2.1) | 1.0 (0.4-1.8) | 0.9 (0.4-1.7) |
| Marker spacing (bp)<sup>7</sup> | 286 (2-1699) | 304 (2-2027) | 324 (2-2173) |

for crosses 3D7xHB3 and HB3xDd2 respectively. Biological replicates were obtained for several progeny clones, where libraries were created from DNA extracted from different cultures of the same parasite clone. These were also sequenced and genotyped separately to enable analysis of concordance between replicates. All sequence data has been deposited at the European Nucleotide Archive and a mapping from clone identifiers to ENA accessions is provided in Table S1 and can alternatively be obtained via the web application at http://www.malariagen.net/apps/pf-crosses where sequence alignments can also be browsed interactively.

Sequence reads from all samples were aligned to the 3D7 reference genome and various metrics were calculated per genome position including depth of coverage and average mapping quality. Visual examination of these data revealed a clear, qualitative difference between a core genome (20.8Mb) comprising regions of near-complete coverage and unambiguous alignments in all samples; hypervariable regions (1.9Mb) where accessibility is severely affected by both extensive paralogy and extreme divergence from the reference genome; and subtelomeric repeat regions (0.6Mb) where accessibility is limited by repetitive sequence (Supplementary Information; Figures S1-S3; Table S2). Hypervariable regions contained all genes in the *var* family, which are known to undergo frequent non-allelic recombination during mitosis (Bopp et al. 2013; Claessens et al. 2014), and almost all genes in the associated *rif* and *stevor* families. Hypervariable regions also corresponded closely with regions of heterochromatin (Flueck et al. 2009) confirming a strong association between chromatin state and qualitative differences in genome variability. All samples exhibited some degree of bias such that coverage was lower where (A+T) content was above 80%, however the high depth of sequencing meant that more than 99.6% of the core genome was covered in all parental clones. Because of the poor accessibility of hypervariable and subtelomeric repeat regions we excluded them from further study and limit ourselves to the core genome for the remainder of this paper. However, raw variant calls and other data for hypervariable regions can be obtained via the accompanying data and web application.

## SNP, INDEL and complex variation within the core genome

SNPs, small INDELs and regions of complex polymorphism were discovered and genotyped within each cross by two independent methods, one based on alignment of sequence reads to the 3D7 reference genome (DePristo et al. 2011; Li and Durbin 2009), the other based on partial assembly of sequence reads and comparison of assembled contigs (Iqbal et al. 2012). Variants where genotype calls in one or more progeny clones were inconsistent with Mendelian segregation (Mendelian errors) were used to calibrate variant filtering for both calling methods (Methods and Supplementary Information; Figures S4-S7). After variants were filtered, both methods achieved near-perfect concordance between biological replicates for both SNPs and INDELs (Table S3) demonstrating that the process from DNA extraction through sequencing and variant calling was highly reproducible. The patterns of inheritance of parental alleles within the progeny of each cross were also highly consistent when comparing SNPs with INDELs (Figure S8) or comparing results of the two variant calling methods (Figures S6 and S7). To provide the greatest possible resolution for the present study, filtered variants called by each method were combined into a single call set for each cross (Table 1; Methods and Supplementary Information). All variant calls can be downloaded from a public FTP site^1^ and browsed via the web application at http://www.malariagen.net/apps/pf-crosses.

The near-perfect reproducibility, concordance between calling methods and highly parsimonious patterns of inheritance and recombination indicate that variants discovered here are true genetic markers, i.e., correspond to genetic differences segregating within the crosses. However, some variants may be false discoveries, in the sense that the variant allele is not correctly ascertained. Also, some genuine variants will have been filtered or not discovered at all. To estimate rates of false discovery and sensitivity we compared variant alleles called in each of the HB3 replicates with the HB3 draft assembly (Birren et al. 2006) and publicly available HB3 gene sequences (Supplementary Information; Table S4). We estimated a SNP FDR between 0.6-2.7% for the alignment-based calling method and between 0.0-1.1% for the assembly-based calling method (Table S5). INDEL FDR estimates were higher for both calling methods, in the range 8.3-12.5%, however we found a high rate of INDEL discordances between previously published sequences (Supplementary Information; Figure S17), making accurate FDR estimation difficult. For sample HB3(1) SNP sensitivity was above 84% and INDEL sensitivity was above 70% for both calling methods, however sensitivity was lower for HB3(2) particularly for the assembly-based calling method (Table S5). This lower sensitivity was partly due to the fact that the assembly-based calling method was only capable of genotyping biallelic variants. All segregating variants in the 3D7xHB3 cross are expected to be biallelic because 3D7 is the reference clone, however segregating variation in the HB3xDd2 and 7G8xGB4 crosses could be multiallelic if each parent has a different non-reference allele at a variant site. This technical limitation accounts at least in part for the lower number of segregating INDELs discovered in the HB3xDd2 and 7G8xGB4 crosses compared with 3D7xHB3 (Table 1).

### INDELs are the most abundant form of polymorphism

Analysis of the combined variant call sets revealed that, within the core genome, segregating INDELs were more abundant than SNPs in all three crosses (Table 1). Overall, 83% of INDELs were found in non-coding regions, where INDELs were 3 times more abundant than SNPs. INDELs were also relatively abundant in coding regions, with the ratio of SNPs to INDELs being approximately 2:1. This relative abundance of INDELs is exceptionally high when compared with other species, for example the SNP to INDEL ratio is approximately 10:1 in primates and 20:1 in bacteria (Chen et al. 2009). The vast majority of INDELs found in the crosses were expansions or contractions of short tandem repeats (STRs), i.e., microsatellites (Figure 1A). In non-coding regions, 83% of INDELs were STR length variations, of which 71% were variations within poly(AT) repeats. In coding regions 77% of INDELs were STR variations, of which the majority were within poly(asparagine) tracts (Figure 1B). Tandem repeat sequences are prone to slipped strand mis-pairing during DNA replication (Li et al. 2002; Lovett 2004) and are known to be associated with high rates of INDEL mutation (Montgomery et al. 2013). Poly(AT) repeats are very common in the non-coding regions of the *P. falciparum* genome (Gardner et al. 2002) and poly(asparagine) repeats are unusually abundant within the exome (Muralidharan and Goldberg 2013; Tan et al. 2010), hence the high INDEL diversity overall may be accounted for by the abundance of STRs in the genome, coupled with the high mutability of tandem repeats due to replication slippage.

**Figure 1.**
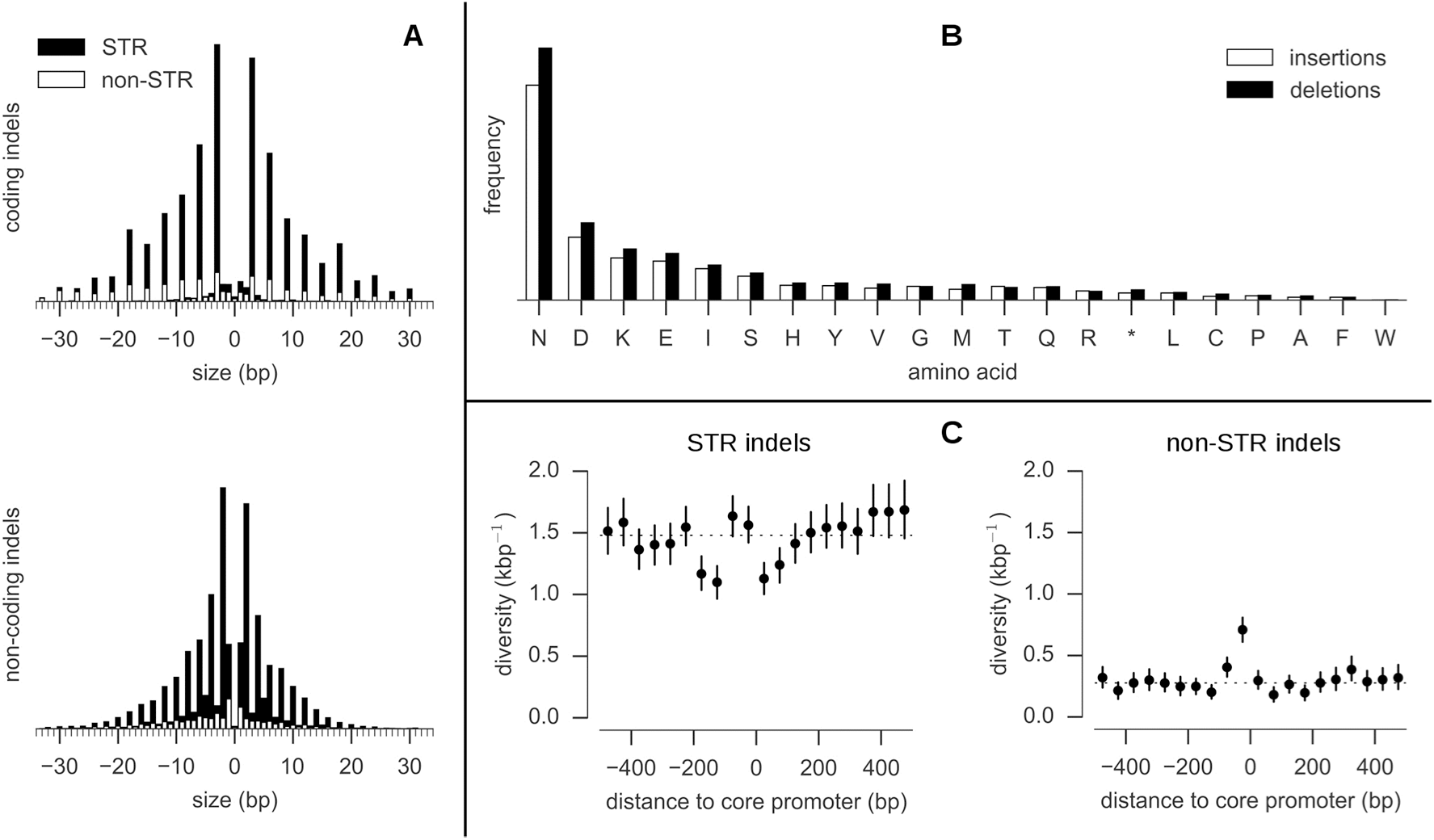
Properties of INDELs. **A**, INDEL size distribution (size > 0 are insertions, size < 0 are deletions). Solid black bars represent the frequency of INDELs that are expansions or contractions of short tandem repeats (STR), solid white bars represent the frequency of non-STR INDELs. Most coding INDELs are size multiples of 3, preserving the reading frame. Most non-coding INDELs are size-multiples of 2, reflecting the abundance of poly(AT) repeats in non-coding regions. **B**, Amino acids inserted and deleted (relative to the 3D7 reference genome). **C**, INDEL diversity in intergenic regions relative to the position of core promoters predicted by Brick et al. (2008). Each point represents the mean INDEL diversity in a 50bp window at a given distance from the centre of a core promoter. Vertical bars represent the 95% confidence interval from 1000 bootstraps. The dashed line is at the mean intergenic diversity for the given INDEL class (STR/non-STR).

Regarding the phenotypic consequences of INDEL variation, frame shift mutations within coding regions are expected to have severe consequences and hence be negatively selected. Here, 94% of coding INDELs were found to be size multiples of 3 and hence preserved the reading frame, whereas most non-coding INDELs were size multiples of 2 reflecting the abundance of poly(AT) repeats (Figure 1A). Within non-coding regions, the potential consequences of INDEL mutations are harder to predict. Relatively little is known about the transcription machinery in *P. falciparum,* however, Brick et al. (2008) predicted the location of core promoters upstream of genes based on a training set of known transcription start sites. Intergenic INDEL diversity in the crosses was found to display a specific architecture relative to the central positions of these predicted promoters, with an excess of non-STR INDELs within the first 50bp upstream of the promoter centre, and a deficit of STR INDELs 100-200bp upstream and 0-100bp downstream of the promoter centre (Figure 1C). INDELs in promoter regions have been shown to alter gene expression in other species (Li et al. 2002) but this remains to be verified in *P. falciparum.*

#### Low nucleotide diversity is punctuated by complex variation in merozoite-stage genes

Average nucleotide diversity across the core genome was 5×10^−4^ per bp in all three crosses (Table 1). However, this relatively low diversity was punctuated by 19 loci with highly diverged alleles, where local diversity over a region up to 2kb was up to 3 orders of magnitude greater (Figure 2). These divergent loci were found almost exclusively within coding regions of genes associated with the merozoite life cycle stage, where the parasite is briefly exposed to the host immune system before invading another erythrocyte, and include several well-studied merozoite surface antigens. The most extreme example was MSP1, a highly expressed protein located at the merozoite surface, where several regions of the gene are known to exhibit deep allelic dimorphism (Ferreira et al. 2003). The complex variation at these loci could not be accessed by the alignment method, because sequences were too diverged from the reference genome, and hence coverage was locally patchy or nonexistent (Figure S9). However, the assembly method was able to construct complete and correct sequences for all parents and progeny in the two main divergent regions of msp1 (blocks 4-11 and blocks 13-16 (Ferreira et al. 2003)) as verified by comparison with publicly available capillary sequence data. Other genes where peaks of diversity were found and alleles could be assembled include four members of the *msp3* family (*msp3, msp6, dblmsp, dblmsp2*), 6 members of the *surf* family (*surf1.2, surf4.1, surf4.2, surf8.2, surf13.1, surf14.1*) and PF3D7_0113800 (encoding a DBL-containing protein with unknown function on chromosome 1). A notable exception to the pattern of merozoite expression was PF3D7_0104100 which is transcribed by the sporozoite specifically within the mosquito salivary gland, suggesting involvement in the early stages of infection (Lasonder et al. 2008). Several of these genes are blood-stage vaccine candidates and/or are being actively studied for their role in erythrocyte invasion, and comprehensive knowledge of variation at these loci is essential for the design of effective vaccines and invasion assays. These results demonstrate that complex variation in these clinically important genes can be ascertained in a high throughput manner via assembly of short sequence reads.

**Figure 2.**
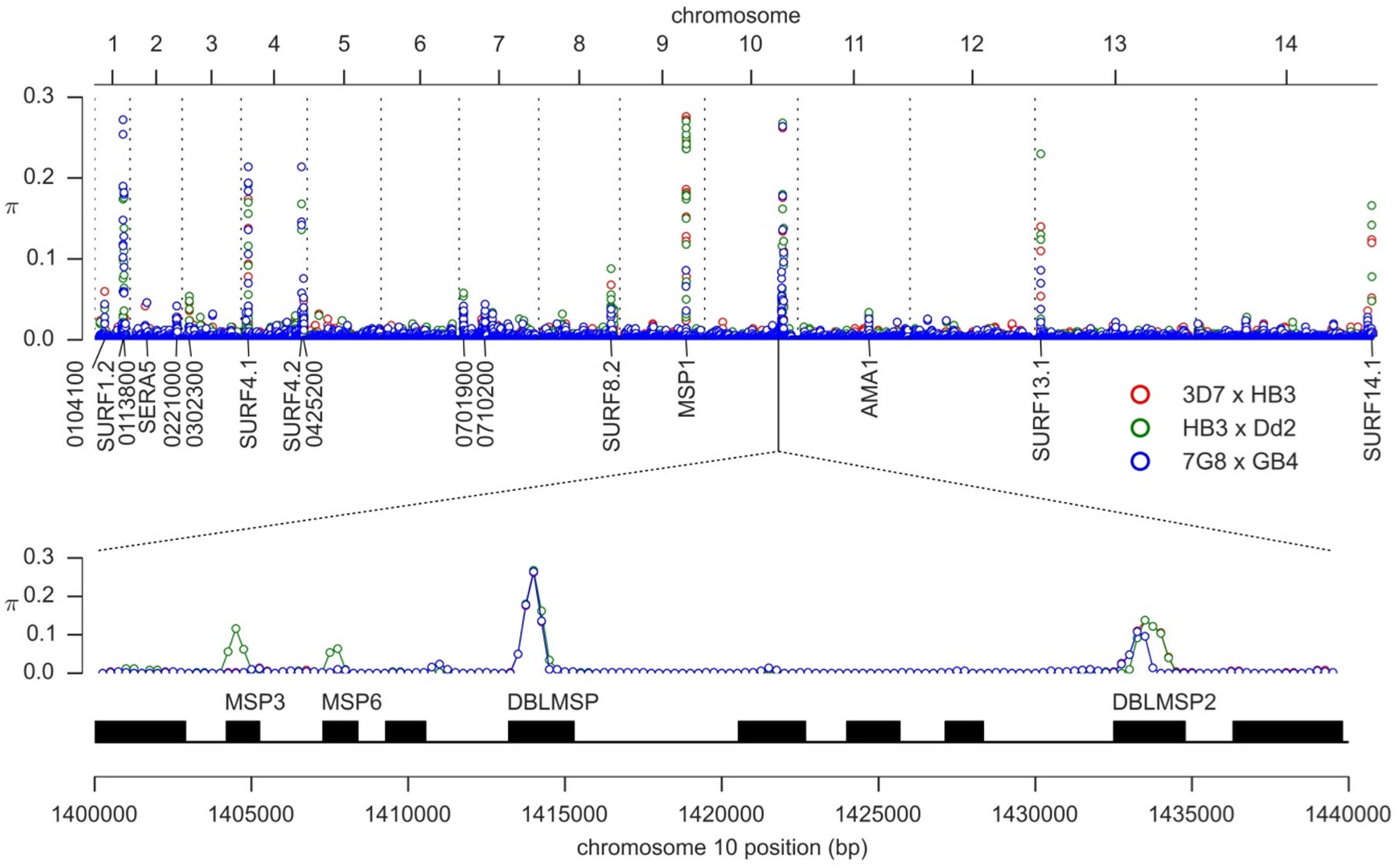
Variation in nucleotide diversity over the core genome. Nucleotide diversity is shown for each cross in 500bp half-overlapping windows across the core genome (which excludes hypervariable regions containing var, rif or stevor genes) using SNPs combined from both variant calling methods and passing all quality filters. The peak of nucleotide diversity in chromosome 10 is expanded to show four distinct peaks due to genes encoding merozoite surface antigens MSP3, MSP6, DBLMSP and DBLMSP2. All labelled loci (with the exception of AMA1) are sites of complex variation where assembly of sequence reads is required to determine the non-reference alleles.

#### Meiotic crossover and non-crossover recombination

These crosses are currently the only available experimental system for *P. falciparum* where meiotic recombination can be directly observed. No previous study has had comparable data for all three crosses, or had the resolution available from deep whole genome sequencing, making this study uniquely able to provide definitive answers regarding the extent and features of meiotic recombination in this species. For each cross, SNP and INDEL variants combined from both calling methods were used as a set of segregating markers for analyses of meiotic recombination (Table 1). The average distance between markers was ~300bp in all three crosses, at least an order of magnitude greater resolution than available previously (Jiang et al. 2011). In eukaryotes, double strand breaks (DSBs) initiated during meiosis are resolved by either crossover (CO) or non-crossover (NCO) between homologous chromosomes (Hastings 1992; Youds and Boulton 2011; Mancera et al. 2008; Baudat and de Massy 2007). A CO is a reciprocal exchange accompanied by a conversion tract, whereas an NCO is a conversion tract without reciprocal exchange (also known as a gene conversion, although NCO events can occur in either coding or non-coding regions; see also Figure 1 in Youds & Boulton (2011)). An algorithm was used to infer CO and NCO events from the size and arrangement of parental haplotype blocks transmitted to the progeny, and to identify both simple conversion tracts (all alleles converted to the same parental haplotype) and complex conversion tracts (containing switches between parental haplotypes) (Methods and Supplementary Information). Because occasional genotyping errors could also manifest as short inheritance blocks, all putative conversion tracts supported by only a single marker or with a minimal length less than 100bp were excluded. This yielded a total of 1194 COs, 230 NCOs and 331 conversion tracts for further analysis (Figures S10 and S11).

#### Gene coding regions are warm-spots and centromeres are cold-spots of CO recombination

Combining CO events from all three crosses, the total map length of the core genome was 15.7 Morgan (95% confidence interval: 14.8-16.6). The total marker span of the physical chromosomes was 21.16Mb giving an average CO recombination rate of 13.5 kb/cM (95% confidence interval: 12.7-14.3). The map length varied between crosses, with 3D7 × HB3 highest (17.7 Morgan), HB3×Dd2 intermediate (16.0 Morgan) and 7G8 × GB4 lowest (14.3 Morgan), although this difference was marginally significant (P=0.06, Kruskal-Wallis H-test) (Figure 3A). There was a strong linear correlation between chromosome size and map length, with 0.55 Morgan predicted for the smallest chromosome (Figure 3B) consistent with ~0.5 Morgan expected if crossovers play an essential role in chromosome segregation and thus the recombination rate is calibrated to produce at least one CO per bivalent (Baudat and de Massy 2007; Martinez-Perez and Colaiácovo 2009; Mancera et al. 2008).

**Figure 3.**
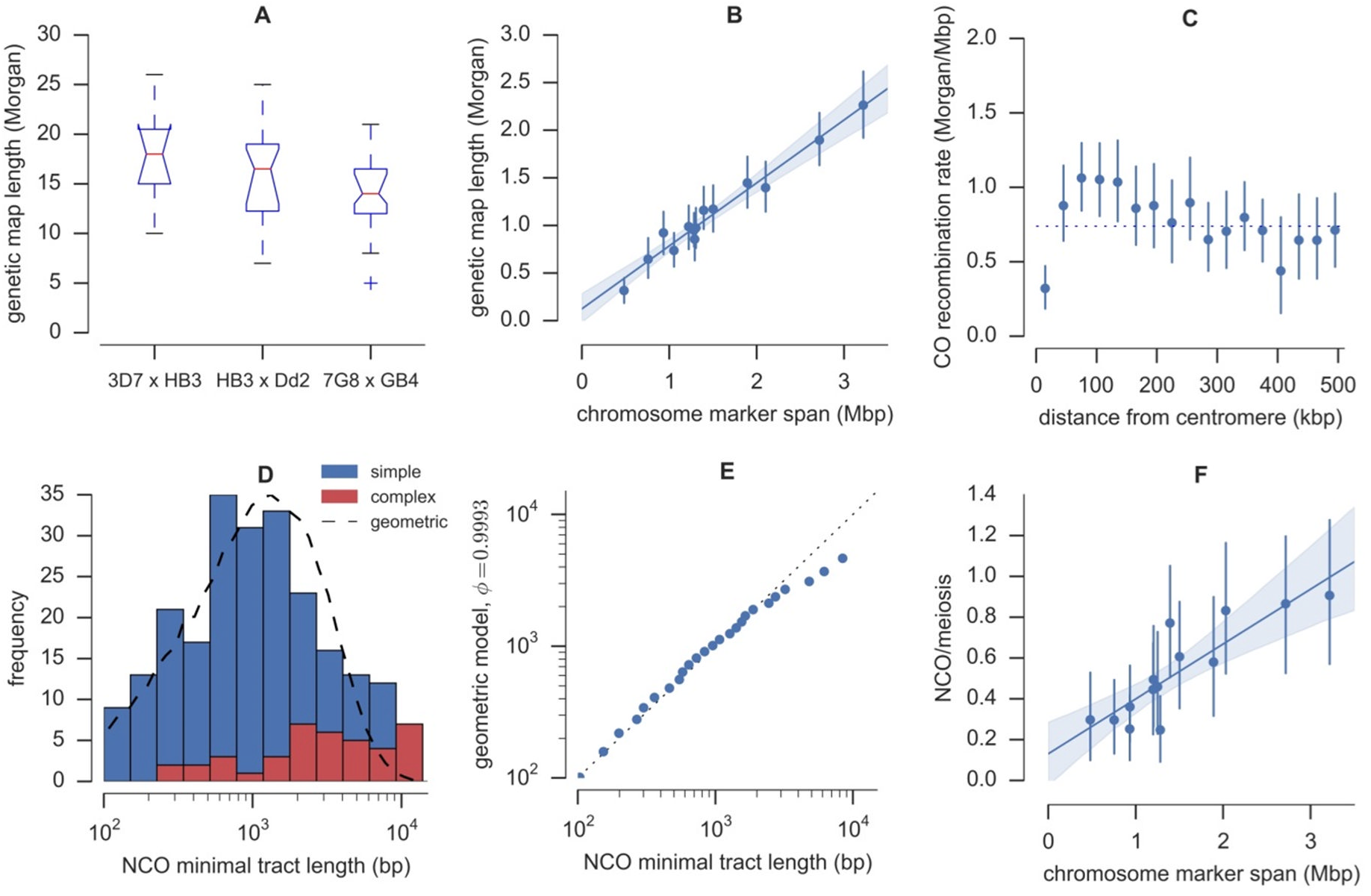
Crossover (CO) and non-crossover (NCO) recombination parameters. **A**, Genetic map length by cross. For each cross the red line shows the median map length averaged over progeny, boxes extend from lower to upper quartiles. **B**, Map length by chromosome. Each point shows the mean map length for a single chromosome averaged over progeny, with an error bar showing the 95% confidence interval from 1000 bootstraps. The line shows a fitted linear regression model with shading showing the 95% bootstrap confidence interval. **C**, CO recombination rate relative to centromere position. Error bars show the 95% confidence interval from 1000 bootstraps. **D**, NCO tract length distribution. The dashed line shows the distribution of minimal tract lengths that would be observed with the available markers if NCO tract lengths follow a geometric distribution with parameter *phi* = 0.9993. **E**, Quantile-quantile plot of actual NCO minimal tract lengths versus the expected distribution of minimal tract lengths that would be observed with the given markers if NCO tract length is modelled as a geometric distribution with parameter *phi* = 0.9993. The data fit the model well except for an excess of tracts with minimal length greater than ~3kb. **F**, NCO frequency by chromosome, adjusted for incomplete discovery of NCO events. Error bars and linear regression as in panel B.

The centromeres were cold-spots of CO recombination, as expected from studies in other eukaryotes and confirming previous data from the 7G8×GB4 cross (Jiang et al. 2011), although the effect was highly localised (Figure 3C). Within ~30kb of the centromere the CO rate was significantly lower, however between ~80-120kb from the centromere the rate was slightly higher than average. Due to the high marker density, in many cases it was possible to resolve the location of CO events relative to individual gene and exon boundaries. Of the 1194 CO events, 396 (33%) were observed within a gene (intragenic COs), 162 (13%) were within an intergenic region, and 636 (53%) were ambiguous (flanking markers spanned a gene boundary). The number of intragenic CO events was significantly higher than expected if CO events were distributed uniformly over the genome (P=0.001 by Monte Carlo simulation). Of the 396 intragenic COs observed, 298 (75%) were observed within an exon, 3 (1%) were within an intron, and 95 (24%) localised across an exon boundary. The number of COs observed within exons was also significantly higher than expected if COs occurred uniformly within genes (P<0.001 by Monte Carlo simulation). Thus a substantial fraction of all CO events were intragenic and occurred within coding regions, in contrast with humans where the majority of recombination occurs within hotspots that preferentially occur near genes but outside of the transcribed domain (Myers et al. 2005).

#### Estimation of conversion tract length and NCO recombination rate

Of the 331 conversion tracts observed, an outlying group of 7 very long (>18kb) complex tracts was found, described further below. Of the remaining 324 tracts, 94 were associated with a CO and 230 were assumed to be NCO conversion tracts. 50% of observed NCO conversion tracts had a minimal size less than 1kb and 73% were smaller than 2kb (Figure 3D). The relatively small size of conversion tracts and the available marker density means that some NCO events will not have been observed, because we required tracts to span at least two markers separated by more than 100bp. To estimate the NCO recombination rate and true tract length distribution, the incomplete discovery of NCO events and bias towards discovery of longer tracts has to be taken into account. In *Drosophila* the distribution of conversion tract lengths was found to fit a geometric model, with parameter *phi* determining the per-base-pair probability of extending a tract (Hilliker et al. 1994). We found that a geometric model also provided a good fit for the observed distribution of tract lengths in the present study with *phi* = 0.9993 corresponding to a mean tract length of 1.4kb, although there was a small excess of tracts observed with minimal length greater than 3kb (Figure 3D, Figure 3E). Assuming this model for the tract length distribution, simulations predicted an NCO discovery rate of 40% for HB3 × Dd2, 39% for 7G8 × GB4, and 45% for 3D7 × HB3 where the marker density was slightly higher (Supplementary Information).

Adjusting for incomplete discovery, the average rate of NCO recombination over all three crosses was estimated at 7.5 NCO/meiosis (0.36 NCO/meiosis/Mb), thus COs are roughly twice as common as NCOs events. The 95% confidence interval for the NCO recombination rate based on sampling error was 6.8-8.1 NCO/meiosis, however this does not account for additional uncertainty in the estimation of NCO discovery rates for each cross. The NCO rate was consistent across chromosomes (Figure 3F) however the correlation was weaker than for the CO recombination rate, presumably due to the fewer number of observed NCO events and thus greater sampling error. As with CO events there was a significant enrichment of NCO events within genes (P=0.002 by Monte Carlo simulation) with 37 (16%) NCO conversion tracts falling entirely within a gene, 110 (48%) spanning a gene boundary, 35 (15%) entirely spanning a gene, and 14 (6%) intergenic.

As mentioned above, 7 apparently long (>18kb) complex conversion tracts were found. Two of these tracts occurred in clone JF6 (7G8 × GB4) within a 60kb region on chromosome 11, and thus appear to be part of a single complex long-range recombination event involving a total of 20 switches in inheritance (Figure S12). Two biological replicates of clone JF6 were sequenced and genotyped in this study, and the pattern of recombination was identical in both replicates. Similar observations were made for clone C04 (3D7 × HB3) where a 70kb region on chromosome 14 accounted for 12 switches in inheritance, and clone 3BD5 (HB3 × Dd2) where an 80kb region on chromosome 10 contained 13 switches (Figure S12). At all of these loci there was no evidence of copy number variation or other artefacts that could manifest as an apparent excess of recombination. These observations do not fit well with conventional DSB repair pathways leading to normal CO and NCO events, suggesting other repair pathways may also be used during meiosis (Mancera et al. 2008) that have more radical results in terms of generating novel haplotypes.

### Recombination within copy number variations spanning drug resistance genes

Clone Dd2 is known to have a three-fold amplification spanning the multi-drug resistance gene *mdr1,* conferring mefloquine resistance (Cowman et al. 1994). Amplifications have also been found spanning *gch1,* conferring resistance to anti-folate drugs, in both HB3 and Dd2, although the amplifications are different in size and extent (Kidgell et al. 2006; Heinberg et al. 2013). The *mdr1* amplification segregates in the progeny of HB3×Dd2 (Wellems et al. 1990) and there is evidence that meiotic recombination has occurred within the amplified region in two progeny clones (Samarakoon et al. 2011a). The *gch1* amplifications have been shown to segregate in the progeny of HB3×Dd2, although one progeny clone (CH3_61) appeared to inherit both parental amplifications superposed (Samarakoon et al. 2011a). Some form of recombination within the amplified region could explain this phenomenon, although the exact nature of the recombination is uncertain. All 5 parental clones have been shown to carry some form of amplification spanning *gch1* (Kidgell et al. 2006; Sepúlveda et al. 2013) thus the sequence data generated in this study provide an opportunity to elaborate on previous results for HB3×Dd2 and extend the analysis of CNV transmission and recombination at drug resistance loci to 3D7×HB3 and 7G8×GB4.

We combined data on depth of sequence coverage and the orientation of aligned read pairs to study CNV alleles in all three crosses (Methods and Supplementary Information). The sequence data confirmed a three-fold amplification in Dd2 spanning *mdr1* and transmission as either 2 or 3 copies to 14 progeny of HB3xDd2 (Figure S13). Evidence for amplifications spanning *gch1* was also clear in all parental clones (Figure 4). The 3D7 reference genome (version 3) has only a single copy of *gch1,* however all studies including ours have found the 3D7 clone to carry multiple copies of *gch1,* suggesting an error in the reference sequence. Parental amplifications spanning *gch1* all differed in extent and copy number, confirming previous findings (Nair et al. 2008; Sepúlveda et al. 2013). We found that alignment of read pairs indicated that the Dd2 amplification was arranged as a tandem inversion (Figure 4D) whereas 3D7, HB3 and 7G8 carried tandem arrays (Figures 4A and 4E) adding further evidence for the independent origin of these CNV alleles. The HB3(2) sample appeared to be a mixture with approximately 20% of parasites retaining the duplication found in HB3(1) and 80% having no amplification (Figure 4D) which is not unexpected given that amplifications can be lost in culture in the absence of drug pressure leading to a mixed colony of parasites (Cowman et al. 1994). Transmission of *gch1* CNV alleles was consistent with Mendelian segregation in the progeny of all three crosses except for two progeny of 3D7×HB3 (C05, C06) and one progeny of HB3×Dd2 (CH3_61) where both parental alleles appeared to be inherited together (Figure 4; Figure S14-S16).

**Figure 4.**
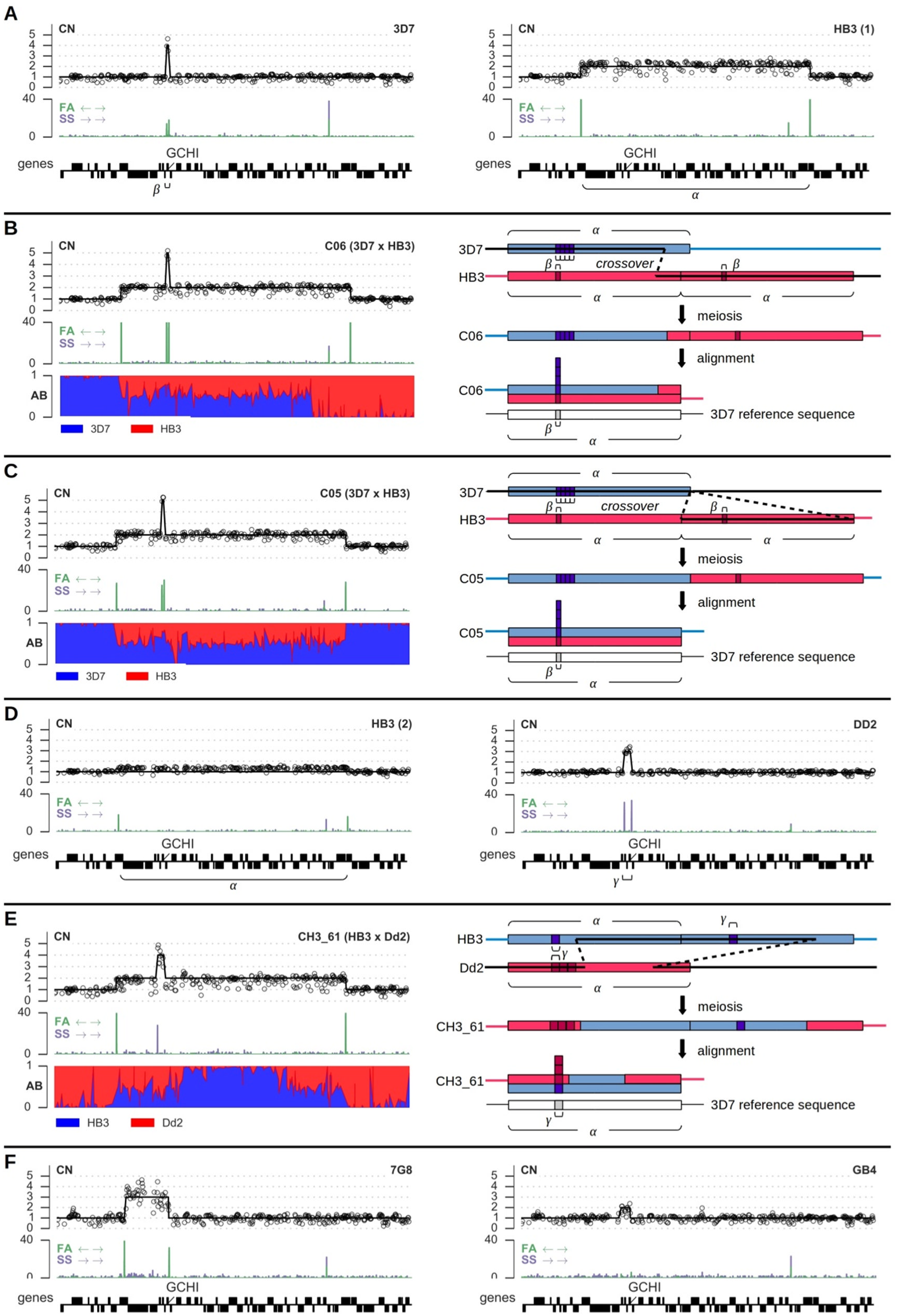
Copy number variation and recombination spanning the anti-folate resistance gene *gch1* on chromosome 12. **A**, CNVs in the 3D7 and HB3(1) parental clones; *α* labels the segment amplified in HB3, *β* labels the segment amplified in 3D7. **B**, CNV and recombination in clone C06, progeny of 3D7×HB3. **C**, CNV and recombination in clone C05, progeny of 3D7×HB3. **D**, CNVs in the HB3(2) and Dd2 parental clones; *γ* labels the segment amplified in Dd2. Note that the HB3(2) clone sequenced here appears to be a mixture, with a minor proportion of parasites carrying the amplification visible in HB3(1). **E**, CNV and recombination in clone CH3_61, progeny of HB3×Dd2. **F**, CNVs in the 7G8 and GB4 parental clones. CN = copy number, markers show normalised read counts within 300bp non-overlapping windows, excluding windows where GC content was below 20%; solid black line is the copy number predicted by fitting a Gaussian hidden Markov model to the coverage data (Supplementary Information). FA = reads aligned facing away from each other (expected at boundaries of a tandem array), SS = reads aligned in the same orientation (expected at boundaries of a tandem inversion), scale is depth of coverage. AB = fraction of aligned reads containing the first parent’s allele.

#### Recombination within amplified regions leads to pseudo-heterozygosity

To further explore the apparent non-Mendelian inheritance of CNV alleles, we considered possible recombination events that could explain the observed patterns of inheritance. Depending how homologous chromosomes align during meiosis, a crossover within a region that is duplicated in one parent could result in a daughter that maintains the same duplication but inherits one copy from either parent for some portion of the amplified region. Within such a segment a haploid individual would become effectively diploid heterozygous for any SNP, INDEL or smaller CNV variants within the segment that segregated between the two parents. We used the alignment of sequence reads from each progeny clone to the 3D7 reference genome to look for evidence of pseudo-heterozygosity and thus recombination within amplified regions. At segregating sites within a region of pseudo-heterozygosity, reads supporting each parental allele should appear in a roughly 1:1 ratio, whereas elsewhere only one parental allele should be observed.

At the *gch1* locus, both clones C05 and C06 inherited the large 161kb duplication from parent HB3 as well as the smaller 2kb 4-fold amplification from parent 3D7 spanning *gch1* only (Figure 4). C06 had a region of heterozygosity spanning the leftmost 130kb of the region duplicated in HB3, but was apparently homozygous for the remainder of this region (Figure 4B). The most parsimonious explanation is that a single crossover occurred within the region duplicated in HB3. Clone C05 had a region of heterozygosity spanning the entire region duplicated in HB3, with borders that appeared to coincide closely with the breakpoints of the duplication (Figure 4C). This is harder to explain, as it would require at least two crossover events at or close to the borders of the duplicated region, which would seem improbable unless the CNV breakpoints are also prone to meiotic crossover (Völker et al. 2010). An alternative explanation is that the HB3 CNV is not a tandem duplication and one copy of the region has been translocated to a different chromosome, however read orientation evidence clearly indicated that the region is tandemly arrayed in both HB3 and C05. For both clones C05 and C06 *gch1* itself lay within the region of effective heterozygosity, thus one copy of *gch1* was inherited from HB3 and 4 copies from 3D7. At the same locus clone CH3_61 inherited the 161kb duplication from HB3 as well as the 5kb 3-fold tandem inversion from Dd2 (Figure 4E). Two separate regions of heterozygosity were visible at either ends of the HB3 duplicated region, which can be explained if two crossover events occurred. Again *gch1* was within the region of heterozygosity, and thus CH3_61 acquired 1 copy from HB3 and 3 copies from Dd2. We also confirmed previous evidence for recombination within the 82kb amplification spanning *mdr1* in two progeny of HB3×Dd2 (Samarakoon et al. 2011a). Clone QC23 had a region of heterozygosity spanning the leftmost 16kb of the segment, and CH3_61 was heterozygous for the rightmost 40kb spanning *mdr1* itself (data not shown). Both of these are consistent with a single crossover having occurred within the amplified region.

### A web application to facilitate data exploration and re-use

The sequence and variation data generated in this study are a rich resource and could serve many purposes beyond the analyses presented here. To facilitate re-use of these data we developed a web application that provides a number of novel tools for intuitive, interactive data exploration, available at http://www.malariagen.net/apps/pf-crosses. The introduction page (Figure 5A) provides navigation to a set of tools, including a tool for browsing and querying a table of variants for each cross and calling method (Figure 5B); a tool for visualising and browsing the genotype calls at individual samples and patterns of inheritance and recombination within each cross (Figure 5C); a tool for browsing the genome, allowing the location of variants to be viewed in the context of genome features and alignment metrics (Figure 5D); and a browser for visualising the sequence alignments themselves, implemented by embedding the LookSeq software (Manske and Kwiatkowski 2009) (Figure 5E). All tools are highly interactive, for example when browsing genotypes the user can hover over any variant and view further information about the reference and alternate alleles, effect prediction, etc. Filters applied to variants can also be changed dynamically, allowing users to explore the entire dataset and compare calling and filtering methods. For the genome browser, a multi-resolution filterbank was implemented to enable highly responsive browsing at all scales from base-pair resolution up to entire chromosomes. The underlying technologies for this web application are being developed as a generic framework so that they can be used with other organisms and datasets, as part of an open source project^1^.

**Figure 5.**
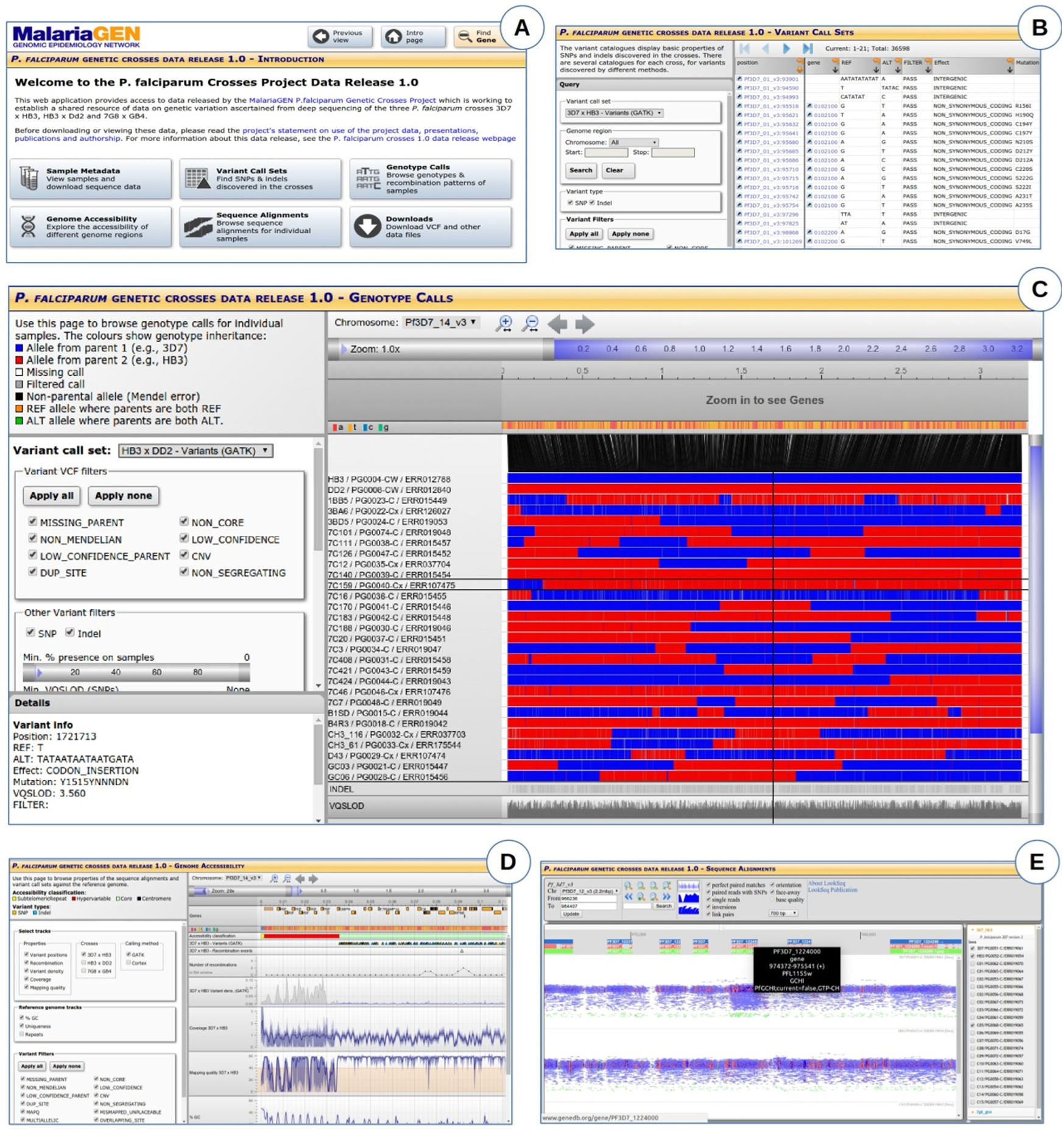
Screenshots from the web application at www.malariagen.net/apps/pf-crosses/ providing access to sequence and variation data on the three crosses. **A**, Introduction page, providing navigation to different tools for data exploration. **B**, Browse and query data on variants (SNPs and INDELs) discovered in the crosses by different calling methods. **C**, Browse genotype calls in parents and progeny and visualise patterns of allelic inheritance and recombination. **D**, Genome browser, providing multi-resolution views of various data tracks including coverage and mapping quality. **E**, Sequence alignment browser (LookSeq).

## Discussion

Here we presented a resource comprising deep sequence data and variant calls for three *P. falciparum* crosses. A fourth cross has recently been performed between the artemisinin resistant clone 803 and the artemisinin sensitive GB4 (Wellems, pers. comm.). The 803×GB4 cross is the last to use a primate host, however new methods are emerging for performing *P. falciparum* crosses using humanized mouse models (Vaughan et al. 2015) opening the possibility for many new crosses to be generated. It would significantly benefit the research community if standardised genomic data on these additional crosses could be generated and incorporated into a single data resource. The web application we have developed for exploring sequence and variation data has a flexible design and could be extended to accommodate further samples and variant call sets, providing a single access point to genomic data on *P. falciparum* genetic crosses.

We have described the first genome-wide data on SNP, INDEL and complex polymorphism in *P. falciparum* spanning both coding and non-coding regions. This does not include hypervariable regions containing *var* genes, because divergence from the reference genome and paralogous sequence present severe challenges to both alignment and assembly-based variant calling methods using short sequence reads. However, we have shown that an assembly-based calling method can ascertain variation in clinically important regions of the core genome where sequences are too diverged from the reference to be aligned, given sufficient homology in the flanking regions. Thus assembly-based methods are clearly capable of dealing with highly divergent sequences and may be able to access some variation in hypervariable regions. However, longer sequence reads will be required to overcome the extensive paralogy, and complete assembly will be required to fully characterise the structural rearrangements between *var* genes that occur frequently during mitosis (Claessens et al. 2014).

The SNP and INDEL variants presented here could be used as a truth set to calibrate variant calling and filtering methods in other studies, for example studies of variation in parasite DNA samples extracted directly from natural infections. By comparing our variant calls with the HB3 draft assembly (Birren et al. 2006) and HB3 gene sequences we estimated that SNP FDR is sufficiently low for this purpose, and INDEL FDR is acceptable albeit higher than has been achieved in some studies of other organisms. The INDEL FDR seems at odds with the fact that inheritance of SNP and INDEL alleles was highly concordant in all three crosses, and INDEL genotypes were almost perfectly reproducible across multiple biological replicates. If we relaxed the FDR matching condition to require only that variants match type and position, estimated INDEL FDR for the alignment-based method was reduced to 5.3-6.2%. The mismatching alleles were always STR INDELs with the correct type (insertion/deletion) and repeat unit (e.g., “AT”) but an incorrect allele length. This could indicate a tendency for the alignment-based method to systematically miscall STR allele length. Some of these mismatches could also be due to genetic variation between HB3 clones with different culturing histories. We also noted considerable discordance between the HB3 draft assembly and published gene sequences regarding INDELs. Although the draft assembly seemed generally more concordant with our variant calls at the 32 genes examined, this was not always the case, and we suspect both the draft assembly and published gene sequences contain INDEL errors. These findings highlight the need for multiple *P. falciparum* genomes fully assembled to the same quality as the current 3D7 reference, so that methods for calling all types of polymorphism and can be accurately evaluated.

We found that INDELs were exceptionally abundant in non-coding regions and displayed a specific pattern of abundance relative to the position of predicted core promoters. Repeat length variations within regulatory regions have been found in other species and shown to affect gene activity (Li et al. 2002; Muzzey et al. 2013). Using the HB3xDd2 cross, Gonzales et al. (2008) showed that both *cis* and *trans* genetic variation influences gene expression in *P. falciparum,* including a major *trans* regulatory hotspot coinciding with the amplification spanning *mdr1.* Variation in gene regulation could affect clinically relevant phenotypes including drug sensitivity, e.g., Mok et al. (2014) found that deletion of a promoter upstream of *pfmrp2* altered sensitivity to quinoline drugs. The data on non-coding variation presented here could provide a starting point for further experimental work to explore the impact of non-coding variation in *P. falciparum.*

*P. falciparum* is a sexually reproducing eukaryotic pathogen, and these crosses provided the first demonstration that parasites undergo meiotic recombination whilst in the mosquito (Walliker et al. 1987). We combined data from all three crosses to estimate a CO recombination rate in the range 12.7-14.3 kb/cM in close agreement with previous studies (Jiang et al. 2011; Ranford-Cartwright and Mwangi 2012). We also estimated that CO events are approximately twice as frequent as NCO events, after adjusting for incomplete discovery of smaller NCO conversion tracts. Samarakoon, Regier, et al. (2011) studied two progeny of HB3×Dd2 using 454 sequencing and observed a similar number CO and putative NCO events in both progeny samples. It is not clear why our estimated NCO rate is lower, especially as marker resolution is an order of magnitude higher in this study and thus power to observe NCO tracts should be higher. We found that conversion tract lengths in *P. falciparum* are comparable to yeast (Mancera et al. 2008) but longer than humans (Jeffreys and May 2004) and *Drosophila* (Hilliker et al. 1994). Our observations of apparent long-range complex recombination events spanning >60kb in some progeny do not fit well with current models for eukaryotic recombination pathways and remain to be explained.

In higher eukaryotes the recombination rate is known to be highly variable over the genome, with most recombination concentrated within narrow hotspots (Myers et al. 2005). Previous work on the 7G8×GB4 cross suggested that the rate of recombination may not be uniform over the *P. falciparum* genome (Jiang et al. 2011). Data from natural populations will be required to robustly evaluate the support for different hotspot models in P. falciparum. We observed a small but significant bias towards recombination within coding regions, however recombination events were frequent in both coding and non-coding regions of the core genome. These different regions have very different nucleotide composition and sequence characteristics, suggesting that a model of highly punctuate recombination targeted at specific sequence motifs is unlikely in *P. falciparum.* Note that these findings apply only to the core genome and entirely different processes may operate within hypervariable regions (Claessens et al. 2014).

We have extended the previous observation of a recombination event within the *gch1* amplification in the HB3×Dd2 cross (Samarakoon et al. 2011a) to illustrate two other cases of recombination within amplifications at this locus. We have also shown that all of these events generate regions of pseudo-heterozygosity within a progeny clone where both parental sequences are inherited and maintained within a single haploid genome. Such events could have important evolutionary consequences. Firstly, drug resistance mutations may confer a fitness cost relative to the wild type allele in the absence of drug pressure (Kondrashov 2012; Anderson et al. 2009; Rosenthal 2013) and may also confer both resistance to one class of drugs and sensitivity to another (Anderson et al. 2009). The process of amplification followed by homologous recombination could provide a mechanism by which both mutant and wild type alleles are acquired and could both be expressed simultaneously, compensating for fitness costs associated with either allele alone. The acquisition of both alleles also creates an opportunity to silence one allele and switch expression between alleles if conditions change. Switching expression between duplicated genes has been shown to occur at the *clag3* locus in response to in vitro drug pressure (Mira-Martínez et al. 2013). Over a longer timescale, gene duplication combined with recombination may facilitate functional diversification, enabling adaptation to different or novel conditions. For example, in plant viruses, gene duplication and recombination may have facilitated adaptation to a wide range of host species (Valli et al. 2007).

Finally, we remark on the connection between INDEL and CNV mutation. Previous studies have found that CNV breakpoints almost invariably occur at sites with some degree of local homology, suggesting that amplifications are due to improper pairing of homologous chromosomes followed by unequal crossover (Nair et al. 2007). Tandem repeats are highly abundant in the *P. falciparum* core genome, thus there are many opportunities for improper pairing during meiosis. Nair et al. also showed that CNV breakpoints are found in repeat regions that are slightly longer than the genome-wide average, thus variations in tandem repeat length could shift the amplification potential to a different set of loci. We have shown here that INDEL variation within tandem repeat regions is abundant throughout the core genome and thus amplification potential is likely to be highly dynamic and variable within natural populations.

The core genome of *P. falciparum* thus appears stable yet poised to undergo rapid evolution within any region that comes under selection. This may become particularly relevant as malaria elimination intensifies in South-East Asia, applying ever stronger selective pressures to parasite populations.

## Methods

### Whole genome sequencing

All sequencing was carried out using Illumina high throughput technology as described in (Manske et al. 2012) except that the PCR-free method of library preparation as described in (Kozarewa et al. 2009) was used.

### Variant calling

Variants were called by two independent methods. The alignment method used BWA (Li and Durbin 2009) to align reads to the 3D7 version 3 reference genome, then applied GATK (McKenna et al. 2010) base quality score recalibration, INDEL realignment, duplicate removal, and performed SNP and INDEL discovery and genotyping across samples within each cross simultaneously using the Unified Genotyper, then used variant quality score recalibration to filter variants, following GATK best practice recommendations (DePristo et al. 2011; Van der Auwera et al. 2013) with some adaptations for *P. falciparum.* The assembly method used Cortex (Iqbal et al. 2012) following the independent workflow. Mendelian errors were used to calibrate variant filtering methods. Filtered variants from both calling methods were then combined into a single set of segregating variation for each cross. Further details are provided in the Supplementary Information.

### Inference of CO and NCO recombination events and conversion tracts

The combined variant call sets were used to infer CO and NCO events and identify conversion tracts within each cross. For each progeny clone, genotype calls were used to identify contiguous regions of the genome where all alleles were inherited from the same parent, by iterating through variants within a chromosome and recording switches in inheritance between adjacent variants. We assumed that any contiguous inheritance block with minimal length shorter than 10kb was not due to a double crossover but represented either the whole or part of a conversion tract (Supplementary Information). Any such blocks occurring in isolation were assumed to be simple conversion tracts. Any such blocks occurring directly adjacent to each other were merged into a single complex conversion tract. To identify CO events, all genotype calls within conversion tracts were first masked, and remaining switches in parental inheritance were called as CO events. Conversion tracts occurring directly adjacent to a CO were then identified, and the remaining conversion tracts were assumed to be NCO events. Putative conversion tracts supported by a single marker or with a minimal length less than 100bp were excluded from further analyses.

### Recombination analyses

To calculate the map length for each cross the identity map function was used because the marker density was high and thus we assumed all crossovers were observed. To estimate the true conversion tract length distribution, conversion tracts were simulated from geometric distributions with different values for the parameter *phi* (the per-base-pair probability of extending a tract). Within each simulation run, the distribution of tract lengths that would be observed given the available markers within each cross was computed. After all simulations were run, the parameter *phi* was fitted to the data by examining quantile-quantile plots comparing simulated and actual distributions of observed tract lengths. These simulations also predicted the fraction of conversion tracts that would be discovered given the markers available in each cross and the requirement that tracts must span at least 100bp. The rate of NCO recombination was then estimated by adjusting the number of observed NCO events by the discovery rate predicted by simulations.

### Copy number variation

The genome was divided into 300bp non-overlapping bins and the number of reads whose alignment started within each bin was calculated for each sample. These binned read counts were then normalised by dividing by the median read count found within the core regions of chromosome 14. Bins where the GC content was lower than 20% were excluded from coverage analyses due to coverage bias in most samples. The fraction of aligned reads with face-away orientation and same-strand orientation was calculated per position for each sample using pysamstats^1^. Copy number state was predicted in all samples by fitting a Gaussian hidden Markov model to the normalised coverage data (Supplementary Information).

### Data access

All sequence data generated in this study are available from the European Nucleotide Archive (ENA). A mapping from clone identifiers to ENA run accessions is given in Table S1. BAM files containing alignments of sequence reads to the 3D7 version 3 reference genome can be downloaded from a public FTP site hosted by the Wellcome Trust Sanger Institute at ftp://ngs.sanger.ac.uk/production/malaria/pf-crosses/1.0/. VCF files containing variant calls for each of the three crosses, from both calling methods as well as the combined call sets, can be downloaded from the same FTP site. All these data can also be explored interactively via the web application at http://www.malariagen.net/apps/pf-crosses/1.0/.

## Acknowledgments

This work was supported in part by the Division of Intramural Research, National Institute of Allergy and Infectious Diseases, National Institutes of Health. Research in LRC’s laboratory was supported by the Wellcome Trust (grant reference number 091791).

## Footnotes

1 ftp://ngs.sanger.ac.uk/production/pf-crosses/1.0/

1 http://github.com/cggh/panoptes

1 https://github.com/alimanfoo/pysamstats

